# Coevolutionary Dynamics of Costly Bonding Ritual and Altruism

**DOI:** 10.1101/060624

**Authors:** Karl Frost

## Abstract

While altruistic behavior and bonding in altruistic pairs or groups of cooperators is observed throughout the animal kingdom, the genetic evolution of such is on an ongoing source of debate, curiosity, and conflict in the behavioral sciences. Many such bonded groups and pairs are observed to take part in costly ritualized movement behavior that is hypothesized to trigger or maintain altruistic sentiments amongst the participants. Such costly ritualized practices could have evolved if they engaged pre-existing behavioral instincts that manifest as altruism in the new context of ritual bonding. While this seems at first to be a ‘Green Beard’ hypothesis (‘marker (ie., ‘green beard’) as honest signal of altruistic intent', an hypothesis well-known to be problematic), it is distinct in two important ways. First, the ritual as marker is costly, and second the ritual engages a pre-existing behavioral potential caused by genes which, importantly, have some other benefit. This paper models the genetic coevolutionary dynamics both analytically and through simulation. It finds that such coevolution can lead to fixation of altruism in a population or to cycling of altruism in the population, depending on the balance of costs and benefits. Where cycling occurs, even though altruism is consistently present in the population, population mean fitness declines with the introduction of these bonding rituals.

## Ritual as reliable signal of altruistic intent

It has been suggested that reciprocal rituals serve to bond humans together into groups and that we share this propensity with many other animals (Rappaport, 1999). Capuchin monkeys will suck each others’ toes, stick each others’ hands in each others’ mouths and stare at each other for minutes at a time, in the context of coalition partnering (Perry et al., 2003). Western grebes (Neuchterlein & Storer, 1982) and great crested grebes (Huxley, 1930) engage in elaborate mutual dances in the context of pair bonding. Cooperative groups have been shown to engage in mutual display behaviors amongst Silver-backed jackal and African hunting dogs (van Lawick-Goodall & van Lawick-Goodall, 1970), pelicans (del Hoyo, Elliot, & Sargatal, 1992), double crested cormorants (Glanville, 1992), and in the pant-hoot dances of chimps (Goodall, 2004; Reynolds, 1965). If rituals successfully bond co-participants into mutually altruistic dyads or groups, they would help groups overcome prisoner’s dilemma type problems: situations where all are better off if all cooperate, but each individual is privately incentivized to not cooperate. Such rituals, taking time and energy which could be spent on other pursuits, are costly. In this paper, I formalize the proposal that some genetically evolved costly ritual behaviors hijack and trigger instincts toward prosociality determined by other pre-existing genes. I then explore the coevolution of the gene frequencies both analytically and through simulations. The analysis demonstrates that in a wide variety of plausible circumstances such rituals lead to the stable evolution of altruism, when such rituals are costly and the genes determining the hijacked behavioral dispositions have independent, pre-existing benefit. As would intuitively be expected, when the pre-existing benefits of the hijacked gene are very high relative to the consequences of cooperation and free-riding, then the hijacked gene remains at fixation and the ritual behavior moves to fixation and, along with it, ritual facilitated altruism. In the case where such pre-existing benefits are of the same order as the consequences of social interaction, then more complex coevolutionary dynamics happen, with mixed populations evolving. Counter-intuitively population mean fitness in this case is reduced despite the increased frequency of altruism in the population.

As background, there are many proposed mechanisms for the maintenance of altruism in groups of unrelated others which have been shown to contextually stabilize or promote altruism. Repeated play and reciprocal altruism can do this (Axelrod & Hamilton, 1981; Trivers, 1971). Cheap punishment has been shown to bolster cooperation and altruism (Henrich & Boyd, 2001). Still, there is the problem of what we do when no one is watching or when punishment can not be dealt out. Neither punishment nor reputation can gain traction when cooperation and defection can’t be observed. In such cases, altruism often decreases, but there are cases where it does not. If there are behaviors that could reliably increase non-enforceable altruism toward non-kin, this would be of enormous benefit to groups. One could speculate that organisms are simplistic and have difficulty making context specific behavior rules and so extrapolate altruistic behavior maintained through such mechanisms as reciprocity and punishment to unwitnessed behavior or to situations which can not be punished or where loss of reciprocal relationship is not a threat. This may be true in some cases (Devetag & Warglien, 2008), but there is no reason to suspect that this would generally be true. The model I present is a novel mechanism which relies on neither context generalization of behavioral rules nor occasional visibility of cooperation/defection.

In addition to demonstrating that such bonding rituals can stably evolve in a population, I demonstrate a number of non-intuitive implications. In cases where the hijacked gene is neutral or non-essential, the ritual may cause the elimination of the hijacked gene through free-rider problems, or the population might have a balance of ritualists and non-ritualists, conditional altruists and non-altruists. In this case, despite the evolution of relatively high levels of altruism and ritual performance, the population mean fitness will decrease. The population goes through damped cycles through the space of gene frequencies. The implication of this is that if the payoff structure is accurately described by this model, any combination of gene frequencies is possible in transition and it is highly unlikely that the population would ever be observed at equilibrium, given the extreme length of time to approach equilibrium.

‘Hijack’ is used here similarly to the concept of sensory manipulation proposed by Dawkins and Krebs (1978), where a behavior is introduced that takes advantage of a pre-existing sensory bias or behavioral tendency to elicit a behavior from the subject. This elicited behavior may or may not be in the subject’s best interest. For example, in guppies, males have evolved orange coloration to take advantage of a pre-existing sensory bias in females to pursue orange things as possible food sources, benefiting the males but disadvantaging the females who would otherwise have a different balance of food vs mate searching (Rodd, Grether, & Baril, 2002). The triggered behavioral tendency is pre-existing and regularly developing. It did not evolve in an environment where it would have been subject to this manipulation and therefore would not be ‘adapted” to it. It is ‘hijacked’ and triggered outside the context in which it had evolved and to which it is adapted, which may then lead to selection pressure against the sensory bias.

The specific kind of hijacking or sensory manipulation referred to here in this ritual model is special. The ritual is intrinsically reciprocal, necessitating the collaboration of two animals to complete. The signal therefore goes to both animals at once. It is simultaneously both a manipulation of the other and a manipulation of the self; it is not possible for one animal to signal the other without themselves being signaled. A synchronous dance, for example, would not be able to be done without a partner to mirror.

It could be that the hijacked gene causes altruistic behavior in specific circumstances where it is warranted for some other reason. The ritual triggers the instinct in a novel context. As an example, humans engage in instinctual ritual activities of mimicry (Weingarten & Chisholm, 2009). Weingarten and Chisholm have suggested that the evolutionary roots of the human instinct to be prosocial toward mimicry partners are in the increased importance of infant-caregiver bonds in an environment of extended childhood development. Mother and child orient on and mimic each other as part of the social transmission of behaviors and have a prosocial reaction in response to this mimicry. However, if some individuals are born that engage in mimicry with others, then they may hijack this mother-child bonding mechanism to provoke altruism in themselves and others in new contexts. When potential cooperative partners meet, they will often subtly mimic each others’ facial and body gestures. When this happens, there is generally a prosocial response, with partners being willing to be altruistic toward each other and thus together they overcome Prisoner’s Dilemma type problems. Of course, a free-rider might then be introduced into the population who can instinctively engage in this mimicry ritual but lacks the mother-child bonding ability and thus the actual prosocial response to the ritual.

Psychopaths have been observed able to strategically engage in these mimicry rituals without having a prosocial response. Of course, more generally and beyond this human example, the original gene would not necessarily code for altruism in the context in which it evolved. It could be a case of pleiotropy, where the gene does not code for altruistic response or helping behavior usually but is ‘tricked’ into it by the ritual.

Humans also engage in socially learned synchrony rituals. These may result in different evolutionary dynamics based in cultural rather than genetic evolution. To avoid confusion, to anchor the imagination in this paper, I will use a toy example based on African wild dogs. African wild dogs are known to engage in elaborate social rituals involving circling movement and vocalizing before a hunt (van Lawick-Goodall & van Lawick-Goodall, 1970). They are also well known for being extremely cooperative and altruistic toward each other, which contributes to very high success rates as hunters. Most likely the actual genetics and history is more complex for actual African wild dogs. The example is meant simply to be illustrative of the model. Imagine an ancestral wild dog species that did not engage in such rituals. It has altruistic cooperative instincts related to mother/pup bonding that involve vocalization and maintenance of proximity. Such instincts are important for the survival and fitness of the individuals. Let us imagine that this altruistic response to vocalization and movement patterns is caused by a single gene. Mother/child aid and cooperation is important for our wild dog’s survival, but there are occasional wild dogs with an alternate allele which have less prosocial response to the proximity and orienting. They do not do as well, but are maintained in the population through mutation. To continue the toy example, a wild dog is born with a mutation in another gene that causes it to try to hyperactively orient on and move with another that is moving with it. When two of these meet, they enter into a feedback with each other that generates a synchronous dance with vocalizations. This dance triggers the prosocial response originally evolved for mother/pup bonding, if the individual has the proper allele of the hijacked gene. The advantage of this is that the pair can take advantage of prisoner’s dilemma type problems in hunting, for example where an individual putting out extra energy in the hunt may not be worth the payoff to itself, but is worth it for the two of them.

If an individual wild dog has the rare non-altruistic allele, then it will enter the physical feedback dance but not be altruistic, potentially free-riding off of altruism of the other. This would then set up a coevolutionary dynamic between these two genes in the population. The trajectory of gene frequencies would be determined by the relative costs and benefits of the ritual, hijacked gene, and prisoner’s dilemma. I put this in mathematical terms below.

## Similar Models

There are two classes of models which are superficially similar to the costly ritual model: green beard models and costly signaling models. It is worth reviewing these before proceeding to mathematically formalize the costly ritual model so that the differences become clear, we avoid conflation, and we can see how different predictions arise.

## Green Beard Models

Where a reliable marker is associated with altruistic intent, positive assortment amongst those visibly marked creates positive assortment of altruists, potentially overcoming free-rider problems. Such markers have been dubbed *green beards*: if altruists and only altruists had green beards, those with green beards could choose to preferentially interact with each other, resulting in groups of only altruists (Hamilton, 1964) (Dawkins, 1976). While such markers have been shown to exist in the world (Gardner & West, 2010), it has been argued that they are likely to be uncommon (Grafen, 1998). Genetic recombination would tend to lead to the breaking of linkages between the genes for the marker and the genes for altruism, and such a very specific pleiotropy, with both caused by the same gene, would be rare. More complex scenarios have been proposed in which green beard markers exist at equilibrium in a population. These rely on some combination of spatial structuring, multiplicity of marker types, or non-existence of pure free-riders (Axelrod, Hammond, & Grafen, 2004; Jansen & van Baalen, 2006; Rousset & Roze, 2007; Traulsen & Nowak, 2007). Interestingly, these models tend not to support a *specific* marker being stable. Instead there is a regular local recurrence of *some* kind of marker associated with altruistic response. Individual markers come and go, as covariance between a marker and an altruistic response arises and then falls apart. What is regularly occurring is not a specific marker being an honest signal of altruism, but that some marker will be an honest signal of altruism. This has been playfully dubbed ‘beard chromodynamics’ (Jansen & van Baalen, 2006).

There are important differences between the payoff structure of the proposed costly ritual model and the green beard model. One difference is that the ritual has a cost where green beard models typically do not have a cost associated with the marker. The cost of the ritual is only paid when another ritualist is encountered. Another difference is that the hijacked gene has some independent benefit. This independent benefit is a plausible assertion, given that something would have facilitated the gene’s establishment in the population in the first place. In order to be a defector, the individual must lack the hijacked gene and incur the cost of this loss. If we reduce both the ritual cost and the benefit of the hijacked gene to zero, we arrive back at the green beard model. As will be shown, in this case the same result is found, that lacking countervailing influences, like spatial structure, such costless rituals hijacking behavioral tendencies produced by otherwise fitness-neutral genes do not lead to altruism at equilibrium.

## Costly Signaling Models

The other class of signal models to which the costly ritual model bears a superficial resemblance are costly signaling models. There is a lot of confusion in the literature and arguably some misunderstandings about the circumstances in which costly signals work and what they predict. In animal behavior literature, Maynard Smith and Harper (2003) differentiate 2 different kinds of costs that are often referred to in signals: *efficacy* costs and *strategic* costs (It is the latter that is the usual subject of costly signaling theory.). An efficacy cost is a cost that is required mechanically in order to make the signal in a way that is perceived and reacted to by the receiver. This is linked to Cronk’s idea of reception psychology (2005), where the idiosyncrasies of the existing evolved psychology or perceptual mechanics requires a cost for transmission. This is exemplified by a ‘manipulative signal’, where a signal takes advantage of a pre-existing perceptual structure or sensory bias of the receiver to manipulate the behavior of the receiver, potentially to the detriment of the receiver (Krebs & Dawkins, 1984; Ryan, 1998). Another example would be an ‘indexical signal’, like stotting in antelope (FitzGibbon & Fanshawe, 1988), where a signal is only possible if one has the quality being signaled. In both of these cases, it may be worth the signaler paying a cost to make the signal. If such a cost is mechanically necessary, it is called an ‘efficacy cost’. If an identical signal that was still indexical were possible without the cost, the costly version would fade out of the population in favor of the costless signal. A *strategic* cost, however, is one where the cost is only possible to pay if one has the quality or a sufficient amount of the quality signaled. This is the usual subject of ‘costly signaling’ models, where the cost itself is necessary for the signal to function (Grafen, 1990; Zahavi, 1975). An individual of lesser quality will find that the cost of the signal is not worth the advantage and so will not pay the cost. In contrast to the case of an efficacy cost, if an identical signal were introduced that was nearly costless, then the individuals lesser in quality would make the signal to get the benefit from the receivers, the receivers would stop paying attention to the signal since it no longer carried useful information, and signaling would disappear from the population.

Key is that such strategic costs do not signal *intent*. They signal *quality,* and *intent* is assumed to be obvious and non-controversial. Costly signaling in this way can not be used to explain the intent to perform altruistic behavior. It may in some cases be a useful explanation of conspicuous altruism itself used *as* a costly signal(Gintis, Smith, & Bowles, 2001), but it is not itself an explanation of a hard-to-fake signal of intent to be altruistic, especially where such altruism is not visible to the recipient.

The literature on ritual behavior amongst humans has generated a range of other theories of ‘costliness’ in relation ritual function, distinct from efficacy and strategic costs. For example, it has been argued that where a belief system changes one’s perspective on and experience of a ritual practice that would otherwise be unpleasant, rituals can be costly-to-fake signals of ‘true belief’ in a group’s belief structure and thus help weed out potential free-riders (Irons, 2001; Sosis & Alcorta, 2003). There has also been a growing literature on the prosocial response to socially learned rituals like synchronous movement (McNeill, 1995). This implies a gene-culture coevolution dynamic. While this is quite similar to the situation modeled in this paper and such a coevolution model has many qualitatively similar results, the descent structure turns out to also have some significantly different implications for the trajectory of behavior in the population over time. These dynamics of socially learned rituals in humans are explored in two separate papers.

## Costly ritual as mutual manipulation: The model and analytic solution for equilibrium

The model presented here is a simple one. Actual evolutionary dynamics are undoubtedly more complex, and single genes coding for complex social behaviors seem improbable. Simple models, however are useful to qualitatively demonstrate evolutionary dynamics (Boyd & Richerson, 1985; Levins, 1966; Winterhalder, 2002), which is the purpose of this paper. In this section, I formalize the verbal model mathematically and explore coevolutionary equilibrium analytically. In the next section, I use simulations to explore the trajectory of the gene frequencies in the population over time

I model the evolutionary dynamics of a reciprocal ritual behavior that hijacks an existing sensory bias to induce altruistic behavior. The ritual is reciprocal in the sense that it involves participation of both individuals and can not be performed unilaterally. A wild dog can not perform synchronous dance and vocalizing unless it has a partner to synchronize with.

Assume a two gene system, each gene with two variants. The first gene codes for the contextual altruistic behavior: allele *A* has this contextual response and allele *a* does not. Allele *A* has a simple additive fitness benefit, g, over allele *a*. In the wild dog example, g is the benefit of better bonding between mother and pup. With all else being equal, *A* would move toward fixation in the population, and *a* would only exist in the population as a vestige or rare mutation.

The second gene codes for ritual performance; allele *E* performs the ritual with willing others (other Es), while allele *e* never performs the ritual. When two individuals meet they are faced with two sequential choices: firstly, whether or not to perform a ritual, at cost f, and subsequently how to behave in a Prisoner’s Dilemma (PD) type situation. The PD has a cost, c, to cooperate. If one cooperates, one’s partner gets benefit, b. f, in the wild dog example, would be the cost of time and energy to engage in synchronized movement and vocalizations. c would be the choice to work extra hard in the hunt or to take greater risks that would benefit others. b would be the benefit to others from these altruistic hunting expenditures. f<(b-c). Cooperation is the altruistic choice. Defection is the selfish choice to not put in any investment that is not immediately of net benefit to the self. The evolutionary equilibrium play for such PD games in the absence of ritual sensory hijacking is to defect.

*Ae* does not perform ritual and defects. *AE* is willing to perform the ritual dance, and cooperates in PD with any ritual co-participant, if they find one. *aE* is willing to perform ritual but defects with all, even with ritual co-participants. *ae* does not perform ritual and likewise defects.

Ae and AE both get the benefit, g, of improved mother/pup bonding. If an AE or aE meets another E, they instinctually perform the energetically costly dance at cost f. An AE performing such a dance will cooperate with their partner, whether AE or aE. aE engages in ritual dance and vocalization behavior but only puts in the minimum necessary in the hunt based on their own personal fitness. aE will defect on their dance partner, but, of course, they also lack the benefit, g, of superior mother/pup bonding. The gene for A/a is invisible, so ritual participants do not ‘know’ a priori if they will receive cooperation or defection. In the absence of E, there is no ritual performance and no cooperation. In the absence of A, there is no cooperation, even if there is ritual performance.

**Table 1:** Encounter outcomes by allele combinations: PD play (Ego/Other) and net fitness benefit (Ego). In PD play, c refers to altruistic cooperation and d refers to defecting/free-riding

|  |  | Other |  |  |  |  |  |  |  |
| --- | --- | --- | --- | --- | --- | --- | --- | --- | --- |
|  |  | AE |  | Ae |  | aE |  | ae |  |
|  |  | PD Play (Ego/Other) | Fitness benefit (Ego) | PD Play (Ego/Other) | Fitness benefit (Ego) | PD Play (Ego/Other) | Fitness benefit (Ego) | PD Play (Ego/Other) | Fitness benefit (Ego) |
| Ego | AE | c/c | $g + (b - c) - f$ | d/d | *g* | c/d | $g - c - f$ | d/d | *G* |
|  | Ae | d/d | *g* | d/d | *g* | d/d | *g* | d/d | *G* |
|  | aE | d/c | *b - f* | d/d | 0 | d/d | -*f* | d/d | 0 |
|  | ae | d/d | 0 | d/d | 0 | d/d | 0 | d/d | 0 |

Table 1 gives the plays in the PD Game and the fitness of allele combinations based on encounters. Table 2 summarizes the fitness equation terms as well as additional parameters for the simulations.

To summarize, population mean fitness values of the different allele combinations, with ω as base fitness are

- V (ae) = ω
- V (Ae) = ω + g
- V (AE) = ω + g + p_AE_b – p_E_c – p_E_f
- V (aE) #x003D; ω + p_AE_b – pEf

… where PY is the fraction of the population with allele Y, and PXY is the fraction of the population with allele combination XY. In the case where there are no E alleles in the population, the population tends to move to PA =1.

Assume haploid genetics with thorough recombination, reproduction in proportion to the reproductive value.

**Table 2:**
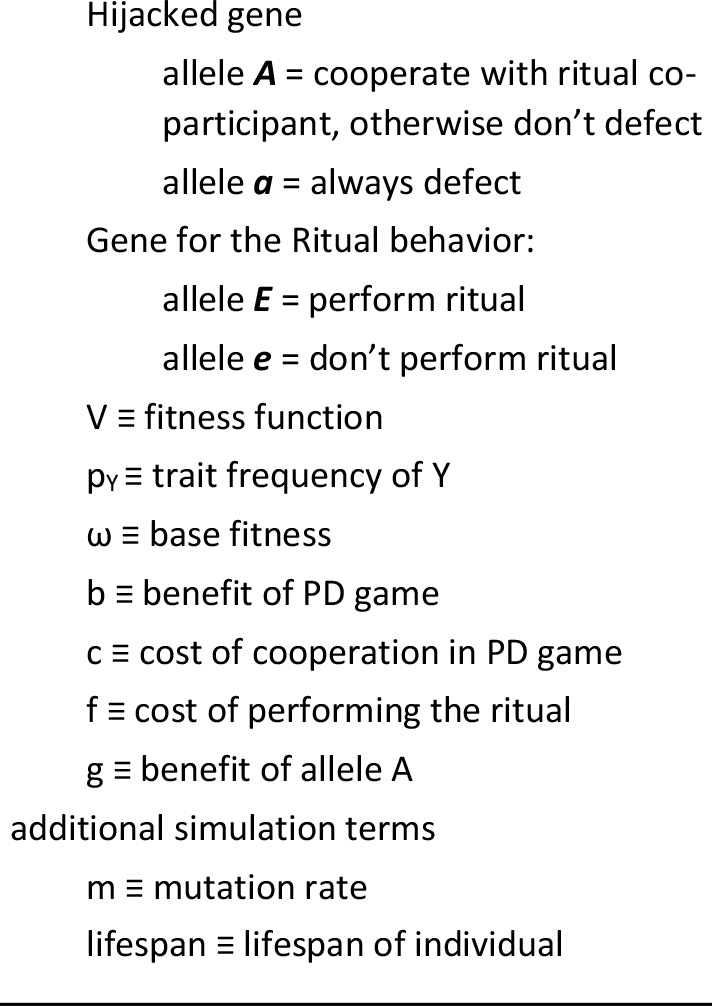
Equation terms

### Results

Doing a little algebra, we find the equilibrium values of pA and pE given in Table 3. The difference in population mean fitness between equilibrium and a population of all Ae is also given. This is taken as the starting population in simulations in the next section.

There are 3 relevant ranges of parameters for the ritual and hijacked allele. For the ritual, we can call these *cheap* (f=0), *costly* (0<f<(b-c)), and *too costly* (f≥(b-c)). For the hijacked allele, we can call these *neutral* (g=0), *useful* (0<g<c), and *very important* (g≥c). There is some parameter combination that would keep any one of the allele combinations at fixation.

The case of *cheap* ritual (f=0) and *neutral* hijacked allele (g=0) matches the conventional green beard model and replicates the results of such. In this case, there is no extra benefit of mother/pup bonding and the ritual triggering the altruistic response is cheaply done. Altruism does not survive in the population at equilibrium, as is the prediction in the simple green beard model. We would expect ritual facilitated altruism to rise sharply at first, but then free riders do very well and eliminate the altruistic allele. Interestingly, the ritual is ubiquitous and thus the potentially altruistic gene is kept out of the population without some countervailing influence. The population mean fitness in the end is unchanged as it rises and then goes back down at equilibrium after the introduction of the ritual gene. Interestingly, the same result is achieved even if the hijacked allele was useful (but not when essential or very important): A is eliminated by ubiquitous cheap rituals. Here, the population mean fitness is reduced by g, the benefit of the hijacked allele. Of course, when the ritual is *too costly*, it fails to evolve. When the hijacked allele is *very useful* it is maintained at fixation in the population, barring rare mutants. In this latter case, if the ritual is *costly* or *cheap* it also moves to fixation in the population as does ritual facilitate altruism, as is reflected in the increase in population mean fitness in the amount of the net benefit of 100% cooperation in the prisoner’s dilemma, minus the cost of the ritual: ΔV = (b−c)−f. This is the scenario that is often envisioned with ritual behavior and bonding and would be reminiscent of the actual populations of African wild dogs, where altruism pays off very well for the group, and ritual behavior and altruism are ubiquitous.

**Table 3:** Equilibrium for different parameter combinations: gene frequencies 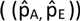and change in population mean fitness from all Ae to equilibrium (ΔV)

| | Very Important Hijacked Allele<br>{$g \geq c$} | | Useful Hijacked Allele<br>{$0 < g < c$} | | Neutral Hijacked Allele<br>{$g=0$} | |
| --- | --- | --- | --- | --- | --- | --- |
| | $(\hat{p}_A, \hat{p}_E)$ | $\Delta V$ | $(\hat{p}_A, \hat{p}_E)$ | $\Delta V$ | $(\hat{p}_A, \hat{p}_E)$ | $\Delta V$ |
| Too Costly Ritual<br>{$f \geq (b-c)$} | (1,0) | 0 | (1,0) | 0 | (0,0)* | 0 |
| Costly Ritual<br>{$0 < f < (b-c)$} | (1,1) | $(b-c) - f$ | $(\frac{f}{(b-c)}, (\frac{g}{c})^{1/2})$ | $-g \frac{(b-c)-f}{(b-c)}$ | (0,0)* | 0 |
| Cheap Ritual<br>{$f = 0$} | (1,1) | $(b-c)$ | (0,1)** | $-g$ | (0,1)**<br>(classic Green Beard result) | 0 |

\*note, analytically, we find for *neutral* hijacked allele that $p_A$ would drift at $p_E=0$ . However, if E is maintained in the population through innovation, then A will be very slightly selected against and $p_A$ moves to 0.
\*\*... similarly here, for *cheap* ritual, $p_E$ would drift at $p_A=0$ . However if A is maintained through mutation in the population, then $p_E$ moves to 1

The case of intermediate parameter values-*costly* rituals and *useful* hijacked alleles (c>g>0 and (b-c)>f>0)-results in a balance of all 4 allele combinations at equilibrium.

- 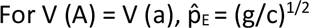
- 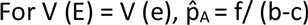

Figures 1A and 1B show these lines of reproductive value equivalence and the direction and magnitude of selection on the traits for different points in trait frequency space for the intermediate case. The equilibrium frequency of the hijacked allele is set by the ratio of the cost of the ritual and the potential net benefit for two altruists in the PD game.

Counterintuitively, this equilibrium frequency of the hijacked allele *does not* depend on how beneficial the hijacked allele is on its own (g), so long as 0>g>c. As the cost of ritual performance (f) increases, the equilibrium frequency of altruistic behavior increases. However, the evolution of the ritual behavior results in a *reduction* of population mean fitness at equilibrium, unless g>c. The *costly/useful* equilibrium is unstable, however. I show in simulations that the population cycles through different frequency combinations, influenced very strongly by the rate of mutation, with a generally declining population mean fitness, from starting point at fixation in Ae. In simulations, starting near equilibrium leads to spiraling through population frequency space away from equilibrium toward a limit cycle made more or less narrow by mutation rate. See Appendix on mutation rate for more details. The selection pressures illustrated in Figure 1 give an intuition for why this cycling would be the case. This decrease in population mean fitness of the population happens despite the evolution of altruism in the population, due to the combined effects of free-riding, reduction of the frequency of the beneficial hijacked allele, and costly investment in ritual.

**Figure 1a.**
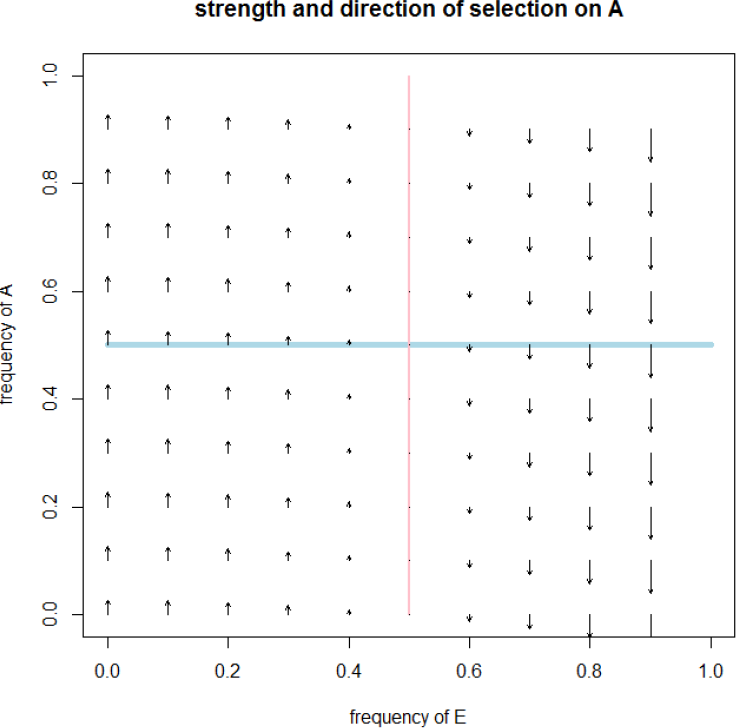
strength of selection on A for intermediate parameter values {*g* = .125, *b* = 1, *c* = .5, *f* = .25}

**Figure 1b.**
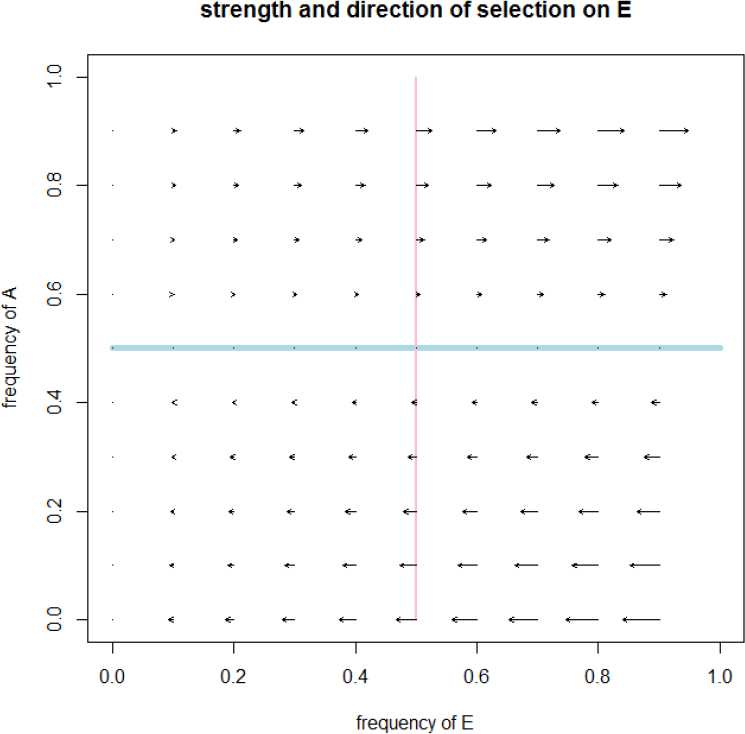
strength of selection on E

These results and the simulations below are based on an assumption of thorough recombination in a well-mixed population. Covariance is assumed to be non-existent. While covariance is often small enough to have negligible effect in evolutionary models, especially in a sexually reproducing population, sometimes even very small amounts of covariance can have dramatic effects. In an appendix, I share analysis assuming zero recombination amongst allele combinations and then again with small, varying amounts of recombination and show that qualitatively similar results are found for the case of zero recombination and that with very small amounts of recombination, the dynamics are almost indistinguishable from the case of complete recombination. See Appendix on recombination for details.

## Simulations

For the simulations, I assume a well-mixed population. Each year, individuals reproduce proportional to the ratio of their fitness to base fitness, divided by their lifespan. Newborns are subject to mutational change from one allele to another (*m*= 2.5 × 10^−5^ changes/generation). After birth/mutation, the population is normalized (to reflect a carrying capacity). See appendix for code. Lifespan (10 years), base fitness (*w* = 10), PD benefit (*b* = 1) and cost (*c* = .5) are held constant amongst simulations. Benefit of hijacked allele (g) and cost of ritual f) are varied to represent the hijacked allele being *very important* (g>c … g =1), *useful* (c>g>0 … g =.125), or *neutral* (g=0), and to represent the ritual being *cheap* (f=0), *costly* ((b−c)>f>o … f = .25), or *too costly* (f>(b-c) … f = 1). The simulations begin with initial conditions, p_Ae_ = 1, representing the introduction of ritual in a population in which the hijacked allele has already evolved to fixation.

**Figure 2:**
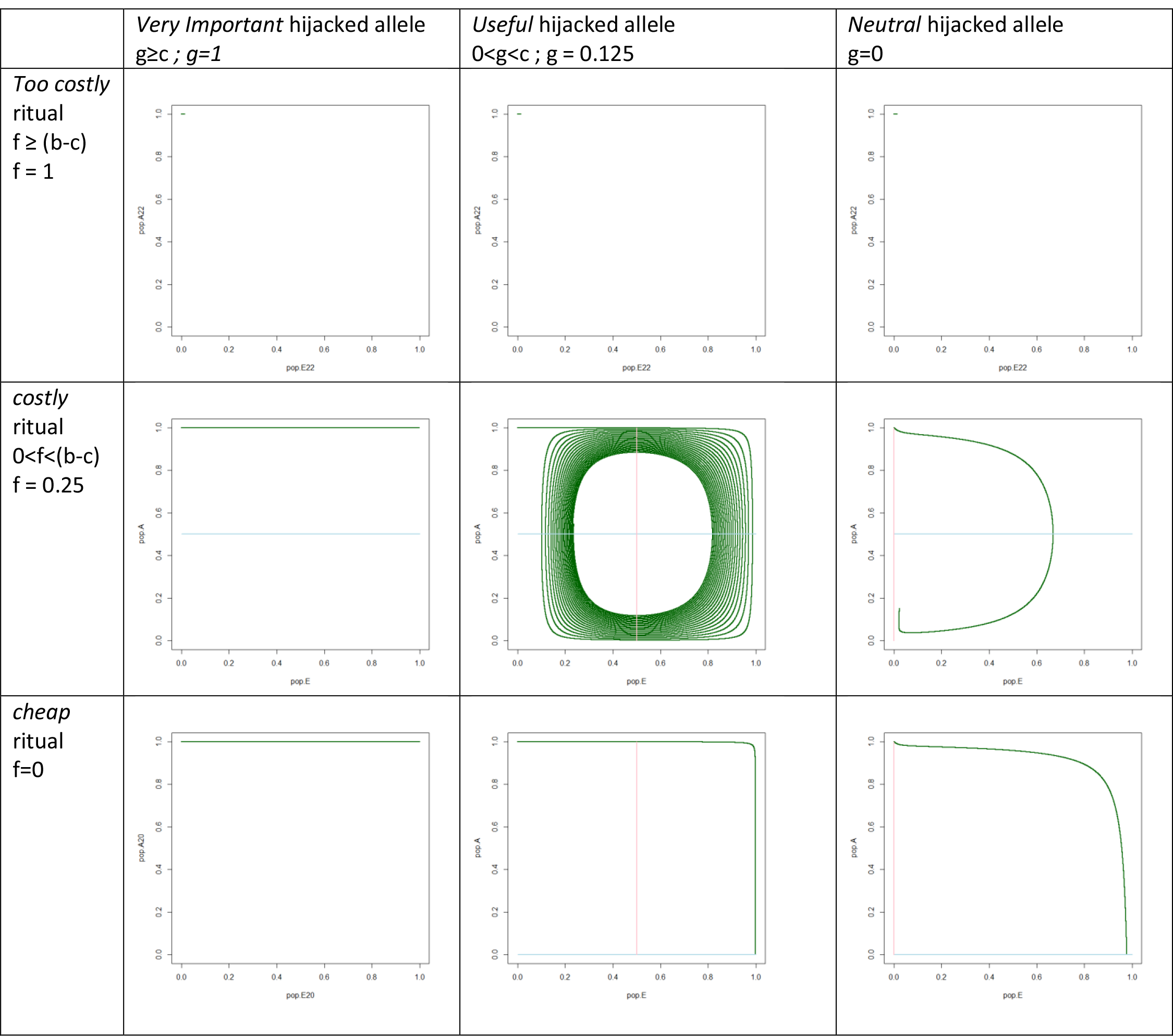
Simulation Results { b= 1, c=.5, w=10, r=.1, m = 1 × 10^−5^) Figure 2A: Gene frequencies

**Figure 2B:**
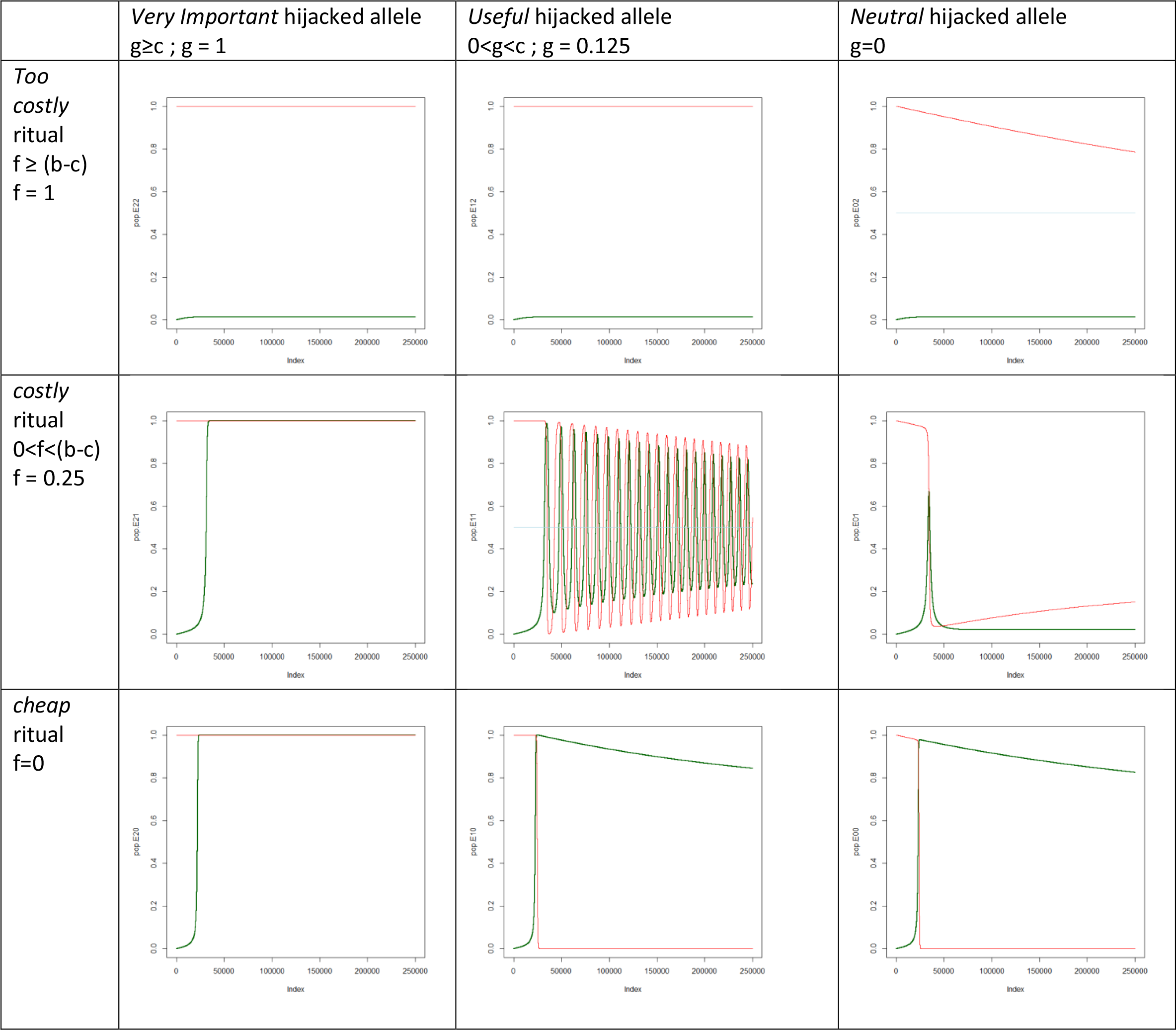
Gene frequencies over time

**Figure 2c:**
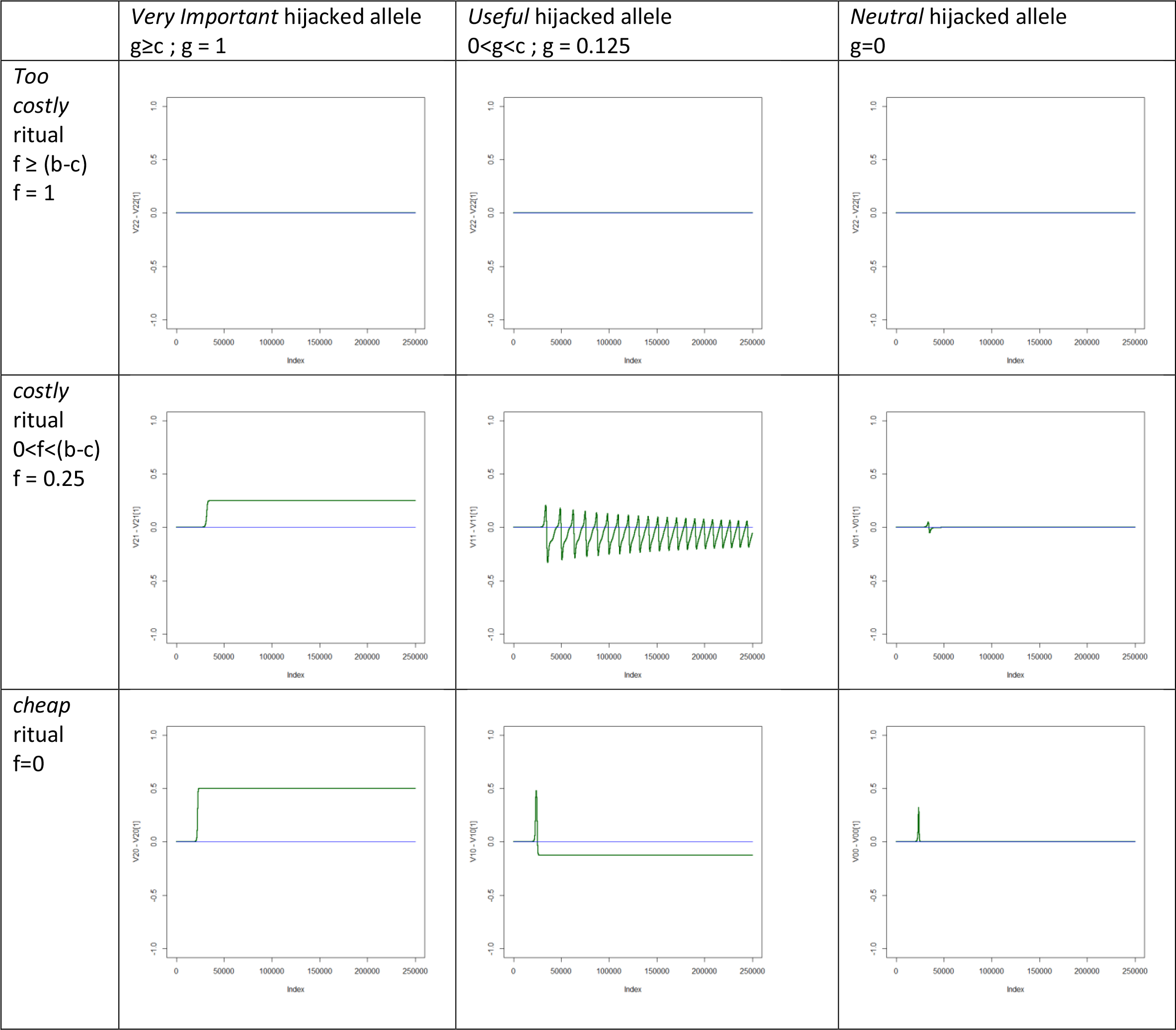
Population mean fitness over time

### Simulation Results

Simulation results are illustrated in Figure 2. Figure 2A shows the trajectory through the space of allele frequencies, Figure 2b shows the allele frequencies over time, and Figure 2C shows the population mean fitness over time. The simulations support the analytic results for equilibrium as well as give a sense of the dynamics over time, which is especially relevant for intermediate values of f and g.

Looking at the effects of the different constants on evolving allele frequencies, for g>c, f<(b-c), the population moves toward fixation in both E and A. The hijacked allele is very important or essential and pre-existing fitness benefit, g, of *A* vs *a* is greater than the potential benefit of free-riding, c. All individuals eventually perform moderately costly or cheap rituals and cooperate with ritual co-participants. Of course, as expected, prohibitively costly rituals (f>(b−c)) do not evolve. For f=0 and g<c, the population moves toward fixation in E and a, which again mirrors the findings for green beards.

For the intermediate values case, where (b−c)>f>0 AND 0<g<c, the population cycles around equilibrium. While the other cases do give fairly rapid evolution toward equilibrium, for such dynamics of *costly* rituals and *useful* hijacked alleles, assuming that a population is at equilibrium is not well justified. The cycling takes place over a very long time scale in comparison to the evolution toward equilibrium in the other cases. For example in this model, in the *cheap* ritual and *very important* hijacked allele system, the population was quite close to equilibrium within 25,000 years. Meanwhile the system with *useful* hijacked allele and *costly* ritual was still in wide looping cycles through frequency space after 250,000 years. The population takes a very long time to even get near equilibrium and en route explores most of the frequency space for the genes. For our wild dog example, in such a case we could find at different moments in time any frequency combination of ritual performers and non, altruists and non. The populations could be anywhere in the space of allele frequencies and would be expected to always be in transition. With smaller mutation rates, the population stays far from equilibrium and moves through a wide limit cycle. The fact that this model generates trajectories through such a wide range of possible trait frequencies points to the fact that simplistic equilibrium-based evolutionary arguments can not be trusted to give reliable predictions for currently observed traits. It points to the necessity for empirical assessment of behavioral dispositions and variance, rather than simple ‘proofs by argument’ about what those dispositions are.

This cycling is very strongly influenced by mutation rate. The simulations were done with a relatively high mutation rate characteristic of mDNA (2.5 × 10^−5^). This causes the population to spiral closer to the (unstable) equilibrium. Lower mutation rates, say on the order of human DNA (2.5 × 10^−8^) result in wide oscillations of the allele frequencies with no time where the population has a significant balance of all 4 alleles: at some point at least one allele is always rare. In summary, these intermediate values cause the population to move through limit cycles. Lower mutation rates cause the limit cycles to be very wide. Higher mutation rates cause the oscillations to be more damped and move the cycling closer to the equilibrium, but not to the equilibrium (see Appendix on mutation rates).

## Discussion and Conclusions

The costly ritual hypothesis has some similarities to Green Beard proposals. The mathematical analysis I present here clarifies the difference between the two and demonstrates how these differences allow for altruism to evolve in a population through novel means that do not rely on such mechanisms as reciprocity, kin selection, or cheap punishment, and which supports altruism in situations where these mechanisms do not. The model is a two gene coevolving system, and for any specific combination of alleles, there is some parameter combination that would support it at fixation. For intermediate values of parameters, there is a balanced equilibrium and the population goes through limit cycles. As explained in the introductory sections, this mechanism for costly rituals to exist in a population should not be confused with arguments about costly strategic signaling of quality, to which there is often an analogy drawn. The different uses of the word ‘cost’ in relationship to signaling have been a source of confusion, so we should be careful not to make unsupportable analogies from one cost and signaling mechanism to another.

While the costly ritual model has some similarities to a Green Beard model, the introduction of cost to the ritual and a secondary benefit to the gene causing the altruistic response creates novel evolutionary dynamics. From the analysis above I highlight the following dynamics.

- If the hijacked trait is essential (g>c), it’s frequency in the population will be unaffected, and costly or cheap rituals ( f<(b−c)) which hijack this gene move to fixation. This results in widespread ritual-facilitated altruism and the solution of PD problems.
- If the ritual is cheap (f=0) and the hijacked allele is neutral (g=0) or is less useful than the potential gains from free-riding (g<c), then ritual performance will sweep to fixation and the hijacked allele will as a result disappear from the population due to the net benefits of free-riding.
- In the intermediate cases, where the ritual is costly (but not too costly) and the hijacked allele is useful, but not essential, then more complex coevolutionary dynamics occur which involve cycling through allele frequencies

The first case is the simplest. A comparison can be made between the ritual model described here and an ‘efficacy cost’ as described by Maynard Smith and Harper. The cost of the ritual would behave identically to an efficacy cost if g were very high – that is, if the hijacked trait was essential. In this case, the ritual would simply serve as a signal to trigger the reaction and the cost would be supported in the same manner as an efficacy cost. Similarly, one would expect a cheaper ritual which could have the same efficacy to do better and so given the possibility, such rituals would evolve toward more cheap rituals. The only differences between such and conventional models of efficacy cost would be that the signaling and the reactions would be intrinsically mutual and that the response would be a specifically altruistic response.

The second case is the parallel to the Green Beard. The costly ritual model is different from the Green beard model, but would be expected to reduce to it in the case of cheap ritual and neutral or non-essential hijacked gene, and this is the result found in the analysis. A long standing view of the problem with green beard effects is that there needs to be some sort of trait linkage for it to work. There is not in this model a linkage between ritual performance and altruistic response to ritual; ritual performance and altruistic response are free to evolve separately. Altruism evolves in a wide range of circumstances with an assumption of complete recombination. There is different trait linkage, however, in the premise that A has both a pre-existing benefit and can be hijacked by a visible behavior (the ritual) to cause cooperation in the PD game. A single gene codes for prosocial reaction to ritual (as in synchronous movement practices) and to some other benefit (like parent child bonding via mimicry), a kind of pleiotropy. One could speculate about the possibility of an evolution of an ability to discern the two situations, to act like A in the original context, but like a in the PD game. Let’s call this allele A’. A’ would in effect allow the breaking of the linkage between prosocial response and the other benefit. The reproductive value of A’ would at all times be greater than that of a and greater than that of A in the presence of ritual and equal to that of A in ritual’s absence. If such a mutation to or modifier of A were feasible, it would dominate and costly ritual performance would rapidly disappear.

This at first seems like an obvious possibility. We can easily see the difference between our mother (with whom we should bond) and strangers (with whom we engage in the PD game). However, we are here modeling the genetically derived, emotion driven heuristics that constrain choices, not our culturally derived ability to distinguish these circumstances. Rather than being as simple as two different ‘person identification’ traits being separated by recombination, this would entail the introduction of a novel trait that suppressed the existing reaction to the ritual in the novel situation. While possible and certainly evolutionarily advantageous in the absence of side effects, the genesis of such a new genetic modifier via mutation is not by any means guaranteed or even expected to be likely, a priori. If it did arise, it would likely come to dominate the population or trigger something analogous to an arms race between the ritual and the hijacked sensory system.

In bearded chromodynamics models, some amount of loose coupling between altruism and marker is necessary to allow altruism in the population. This covariance in these models is the result of sochasticity. Costly ritual dynamics are different in that such coupling is unnecessary for the evolution of altruism. The models here assume no covariance and still altruism is able to evolve, not only in the case of an essential hijacked gene, but also in the intermediate case (0<g<c and 0<f<(b−c)). The same applies when covariance is allowed, as demonstrated in the Appendix. However, in the intermediate case for costly rituals, there is cycling of the population through frequency space, with the oscillations only damping toward equilibrium under strong mutational pressures.

Because of the cycling dynamics potentially caused by this payoff structure in the intermediate case, the population may never have time to get near equilibrium and thus may at any one given point be found anywhere in the space of gene frequencies. In the cases of very useful hijacked trait or cheap ritual, coevolution is rather fast and direct toward equilibrium. We are then safer in the conventional assumption that the population is near equilibrium, and equilibrium analysis may give us a useful prediction about expected gene frequencies. The intermediate case, however, presents us with a problem. The population moves through damped oscialltions down to a limit cycle over a very long period of time, potentially exploring all of the space of gene frequencies in the process. Simple equilibrium analysis may not give us a very useful prediction, then, about gene frequencies. Given the length of time to get near to equilibrium and the range of allele combinations explored on the way there, there is no good justification for assuming that the population would be near equilibrium. We don’t have a way to predict, a priori, what the gene frequencies should be, merely how the system might evolve over time, if we know the gene frequencies and the parameters. Moreover, it is also quite interesting that in this intermediate case, the introduction of ritual behavior will lead to a *decrease* in the average fitness of the population over time, despite the evolution of altruistic cooperation in the population. Unlike bearded chromodynamics or costly signaling models, ritual hijacking does not in the long run benefit the overall population *unless* the hijacked gene is sufficiently vital.

While there are costs involved and they are necessary for the evolution of altruism with in the costly ritual model, the mechanism for the stabilization of altruism is quite different from the evolutionary mechanisms that stabilize strategic costly signals of quality. Further, the cost of the ritual facilitates the evolution of honesty of signal of altruistic *intent*, rather than a guarantee of some level of *quality* in a context of guaranteed intent. Findings from models of costly signaling of quality can not be extrapolated to situations where intent is being communicated.

With this model, I’ve shown that costly rituals can evolve in a population. What happens if the ritual *cost* were itself allowed to evolve? Would higher or lower cost rituals dominate? Would ritual cost increase or decrease, or would there evolve a range of ritual costs? These simple questions could be taken a variety of ways. Assume that, in addition to the non-ritualist allele, there are two different ritual alleles in the population, coding for ritual performance with costs f and f’, f>f’. It might be that the two ritualist types recognize each other and do some form of ritual together, or they might not recognize each other. If they recognize each other, do they default to the more or less costly ritual? Do they each perform their own ritual, and is this difference visible or not to the other? Do the two rituals have the same effect on Prisoner’s Dilemma play (b=b’, c=c’) or is PD play proportional to f (f’/f = c’/c= b’/b)?

These questions are explored in an appendix. In a simple case, where the two rituals have the same result (b=b’, c=c’) and are mutually recognized then, trivially, the less costly ritual eliminates the more costly, as it always has higher fitness than the more costly one. This then also means that pa is reduced with the introduction of the cheaper ritual, it being proportional to f. However, if PD investment is *proportional* to ritual investment and they recognize each other and play the same when different types meet (either low or high), then the two can coexist in equilibrium, unless g>c, in which case the *higher* investment ritual wins. Many different outcomes are possible, ranging from one or the other type dominating to coexistence, depending on how the cheaper ritual relates to PD play, how the two types act when they meet each other, and parameter values. For more details, see Appendix on multiple rituals of different costs.

This theoretical model offers a number of implications for empirical research. First, it shows that such ritual behaviors can evolve which facilitate altruism in situations that can not be explained through reciprocity, kin selection, or the possibility of cheap punishment of non-altruists. This would suggest that such costly rituals would be more likely to occur where there are more significant gains to be had from altruistic cooperation and also that they should be able to still solve PD problems where surveillance is not possible (and reciprocity or punishment can get no traction) and individuals are less related (and thus kin selection would be less of an issue). With hunting relations, this could mean more voluntary sharing or risk taking. With mating pairs, it could mean greater monogamous exclusivity. In examining animal ritual behavior, it would be most interesting to look for such altruistic responses that can not be explained through these other mechanisms (reciprocity, cheap punishment, kin selection).

If the benefits of the hijacked allele are less than the net benefit of free-riding and the cost of the ritual is less than the benefit of reciprocated altruism, the model predicts mixed populations of ritual performers and non-performers. In such cases, there should at all times be some genes under recent selection pressure, as the population moves through cycles, whether the genes for ritual performance and any genes for altruistic response. If instead the hijacked allele were more essential, the populations would be expected to move more rapidly to fixation. If genes responsible for these behaviors could be identified, this represents another testable hypothesis of the model.

Of course, this is a purely genetic model and may not describe accurately the evolutionary dynamics of socially learned rituals. We would expect much faster evolution of socially learned synchrony rituals like coordinated dance or close order drills (McNeill, 1995) due to the relative speed of cultural evolution processes (Perreault, 2012). Where there are coevolutionary dynamics between a socially learned ritual form and hijacked genetically determined predisposition to cooperate with synchrony partners, we would also expect much more rapid genetic evolution, based on fast culture-led gene culture coevolution, a question explored in a separate paper.

## Appendix 1: R code gene-gene coevolution, no stochasticity

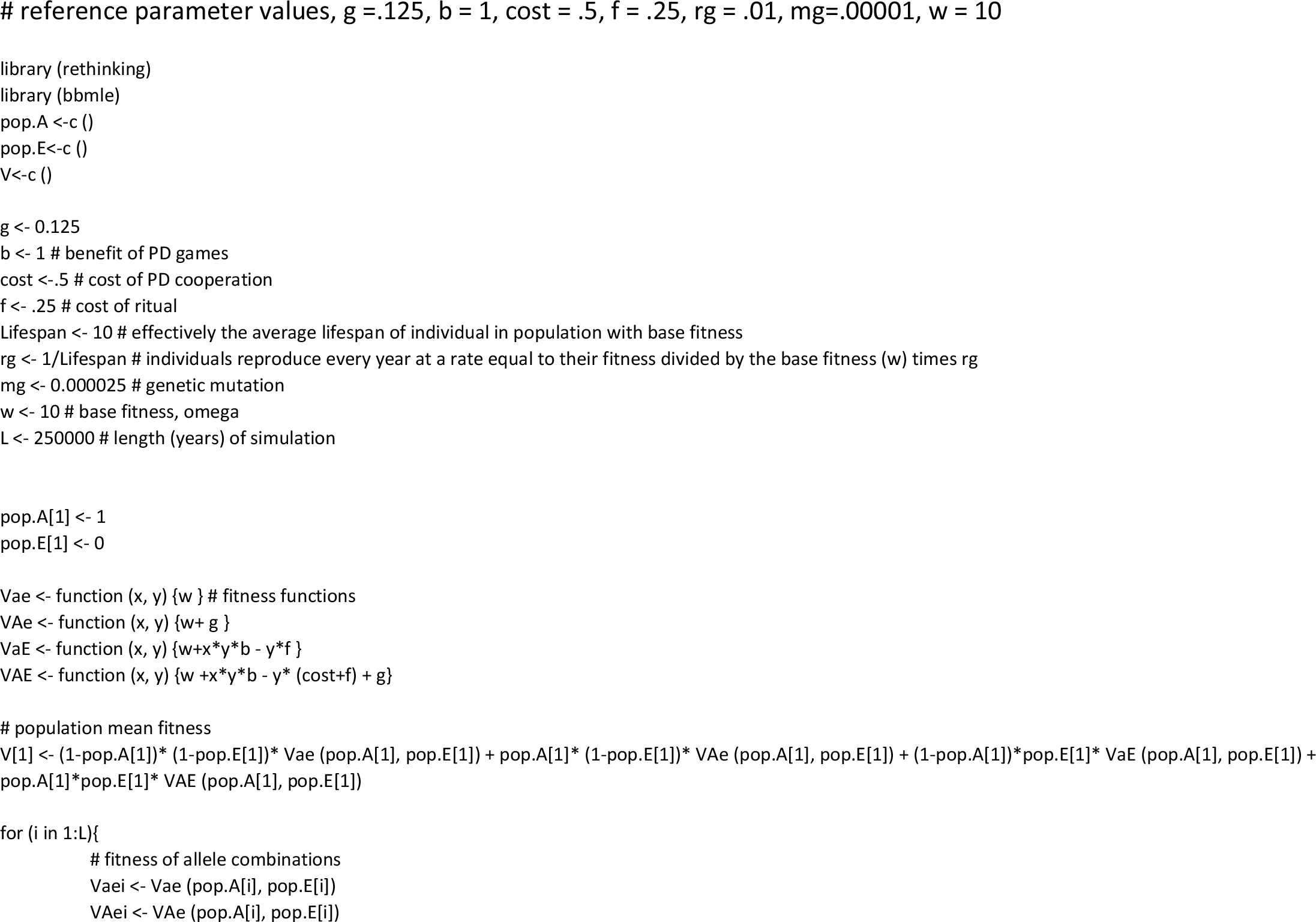
R code gene-gene coevolution, no stochasticity

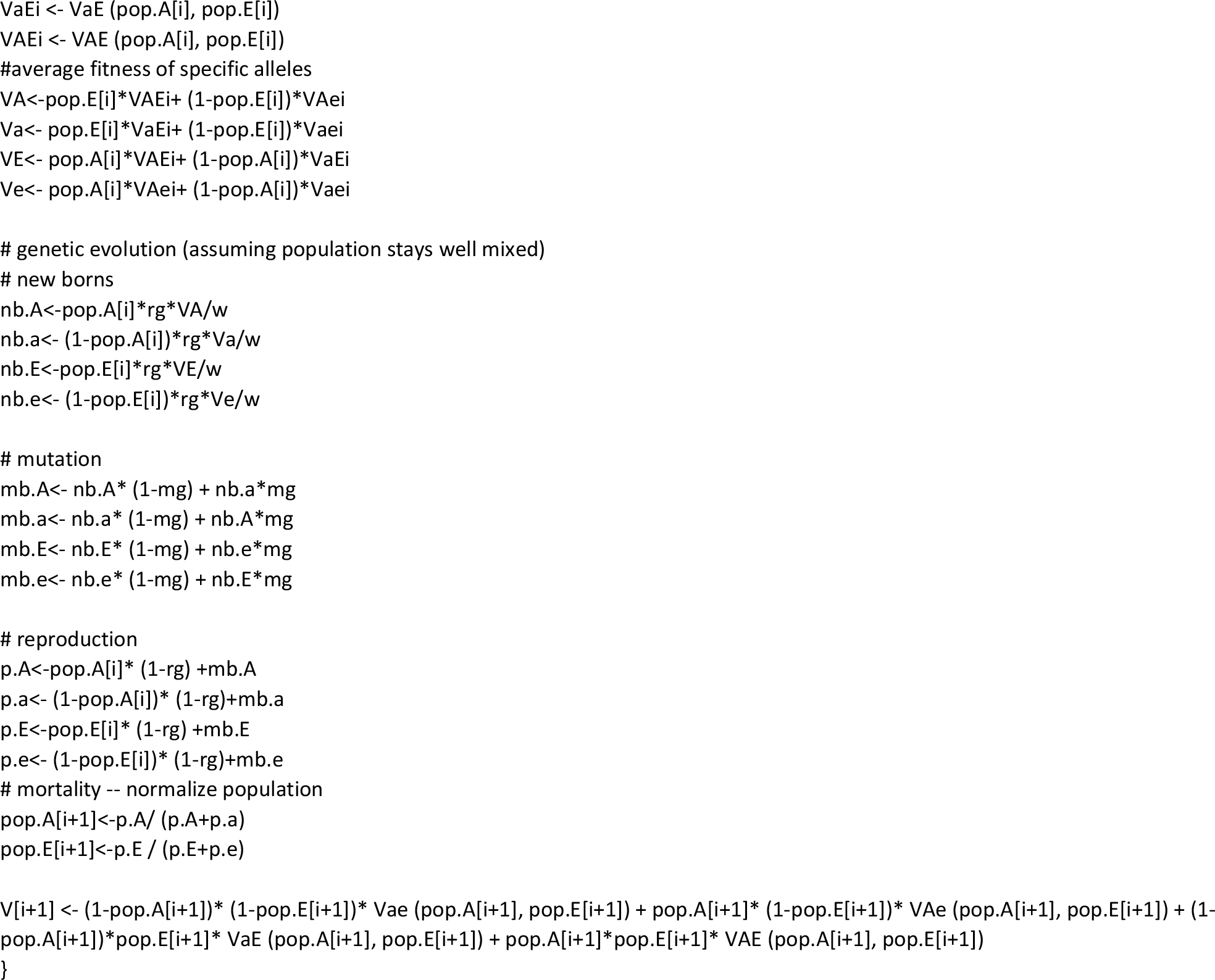

## Appendix 2: the effects of mutation rate

For (b−c)>f>0 AND 0<g<c, the population cycles around equilibrium. It turns out that this is very strongly influenced by mutation rate. Often times simulations are done with little thought communicated about why a particular mutation rate is chosen or even without concern about whether it is realistic or not. While this may be of little consequence in many evolutionary systems, in others this seemingly innocuous assumption may have extraordinarily strong and less-than-intuitive impacts.

Mutation rates for DNA vary tremendously. On the relatively high end for animals, there are mitochondrial DNA mutation rates: 2.5 × 10^−5^ changes per location per generation. On the upper end (at least for humans, a particularly well-studied organism), DNA with proof-reading and self-correction capacity can have a much lower mutation rate, on the order of 2.5 × 10^−8^ changes per location per generation.

A relatively high mutation rate characteristic of mDNA causes the population to spiral farther in toward the equilibrium. Lower mutation rates, say on the order of human DNA, result in oscillations of the allele frequencies with the population at no time having a significant balance of all 4 alleles: at some point at least one allele is always rare. The Figure below demonstrates the results of simulations for high and low realistic mutation rates and for a population beginning with either fixation of the hijacked allele and no ritualists or near the equilibrium.

It should be noted that mutation is just one way to introduce rare alleles and that others might exist and have stronger effects, like migration from partially isolated communities in a different environment.

**Figure X:**
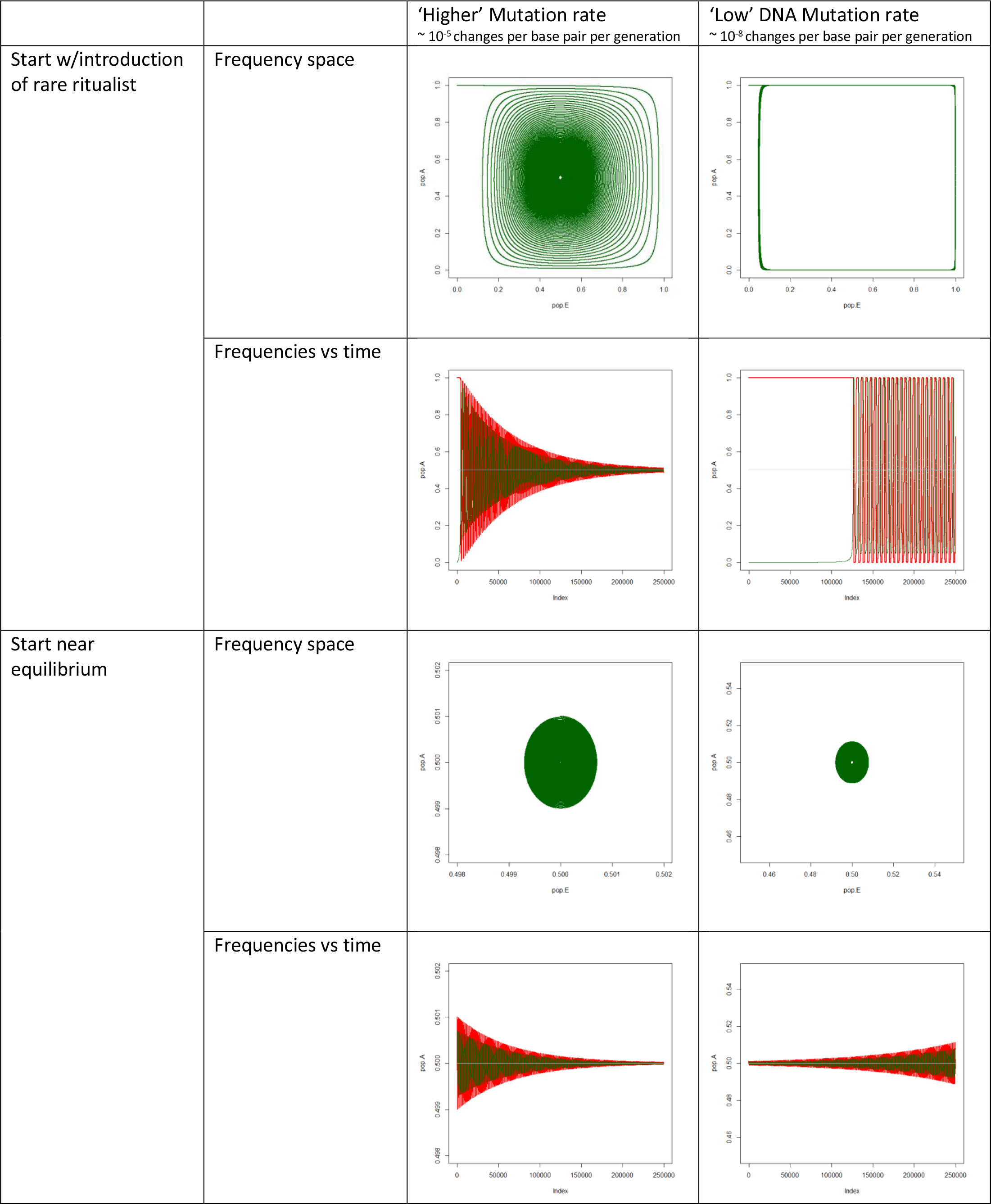
The effects of mutation rate on trajectory of allele frequencies for intermediate cost of ritual and benefit of hijacked allele, fast reproducing species (r =.5)

## Appendix 3: Variance in costliness of ritual

A very reasonable question to ask might be if a ritual that is otherwise stable in a population is stable against the introduction of either a less or more costly ritual. Of course, the cost of the ritual might be a requirement of the sensory system, a necessary efficacy cost. In this case, it would not be necessary for such a ritual to be stable against the introduction of a less costly ritual, since such would be impossible. For example, engaging in a mimicry takes some minimum amount of time and energy to establish the “I know that you know that I know that you know” of the mutuality. It may be that there are different versions possible that may be more or less costly, but some minimum investment might reasonably be assumed.

Given this range, will a system with two different rituals (E and E’) of different costs (f= h*f’, h>1) evolve toward the lower or higher cost ritual at fixation or dominating the other but in balance with non-ritual performance, or would all 3 alleles coexist in balance at equilibrium? It turns out that outcomes very much depend on the way that the two rituals relate to each other and how they relate to the play in the prisoner’s dilemma game. For the latter, we might consider variants where they play the same (b=b’, c=c’) or where play is proportional to ritual investment (f/f’ = c/c’ = b/b’). Play being proportional to ritual investment might mean that the extent of altruism is depending on how much visible investment there is in the ritual. Of course, for both rituals a allele individuals do not pay the cost c of cooperation and contribute nothing in the context of the PD game.

The following was found through equilibrium analysis. In the case of a balance at equilibrium, this may take an enormously long time to reach and move through a wide range of frequency combinations enroute, or it may indicate a balance point around which the population cycles indefinitely. Simulation results in Figure A3 suggest pathways of the populations over time for specific forms of relationship between low and high cost rituals.

## Mutually unrecognized, play same

If they are mutually unrecognizable (treat each other as non-ritualists), there would be a potential path dependency. If one of the rituals were to be at fixation or close and moving toward it rather than cycling, then it would crowd out the other ritual, irrespective of which is higher or lower cost. The one that gets to fixation first has all the cooperative partners and crowds out the other ritual. However, the cycling of the populations allows ample opportunities for competition if this is not the case.
If they are mutually unrecognized, play the same PD plays, and do not go to fixation, then the lower cost wins. (See Figure 3A for simulation)

## Mutually unrecognized, play proportional to ritual investment

If play is proportional to ritual investment, then a bit of algebra shows that there is a shared equilibrium. For h= f’/f, PA = f/ (b−c), P_E_ = sqrt (gh/ (gh+c)), P_E’_ = P_E_ /h. The less costly ritual does better, but they coexist. (See Figure 3A for simulation)

## Mutually recognized, play same PD plays (f>f’, c=c’, b=b’)

If they are mutually recognizable and play PD games the same, then the lower cost ritual wipes out the other; it always has an equal or higher fitness than the higher cost ritual play. The long term stability of costly rituals may depend on a mutual recognition of invested cost in the ritual. If a player evolves that can do a ritual at lower cost that triggers the same response in the hijacked allele and is mutually recognized by higher cost ritualists, then such lower cost ritualists would take over the population, though perhaps remain in balance with non-ritualists for g<c

## Mutually recognized, if mixed pair, default to invest in ritual and play lower amount (f/f’ = c/c’ = b/b’)

In the case that g>c, the higher cost ritual moves rapidly to fixation. E individuals will always do at least as well as E’ individuals who always do better than e individuals. If g<c, then there is an internal equilibrium. It is a very messy expression, but from simulations, it can be characterized that the equilibrium of the higher cost ritual is slightly reduced by the presence of the lower cost ritual and the combined frequencies of both rituals is a little lower than what the lower cost ritual would achieve on its own.

## Mutually recognized, if mixed pair, default invest and play higher (f/f’= c/c’ = b/b’)

The equilibrium is the similar to default lower. Similarly if the higher cost ritual moves to fixation for g>c, it wipes out the lower cost ritual, not by crowding out, but because of the lesser degree of altruism amongst low cost ritualists when they encounter each other. They would be able to share in altruism of high investment rituals when at low frequency but become progressively more disadvantaged as they get more do worse when they encounter each other. For g<c, the two alleles exist in balance. See simulation results in Figure A3.

**Figure A3:**
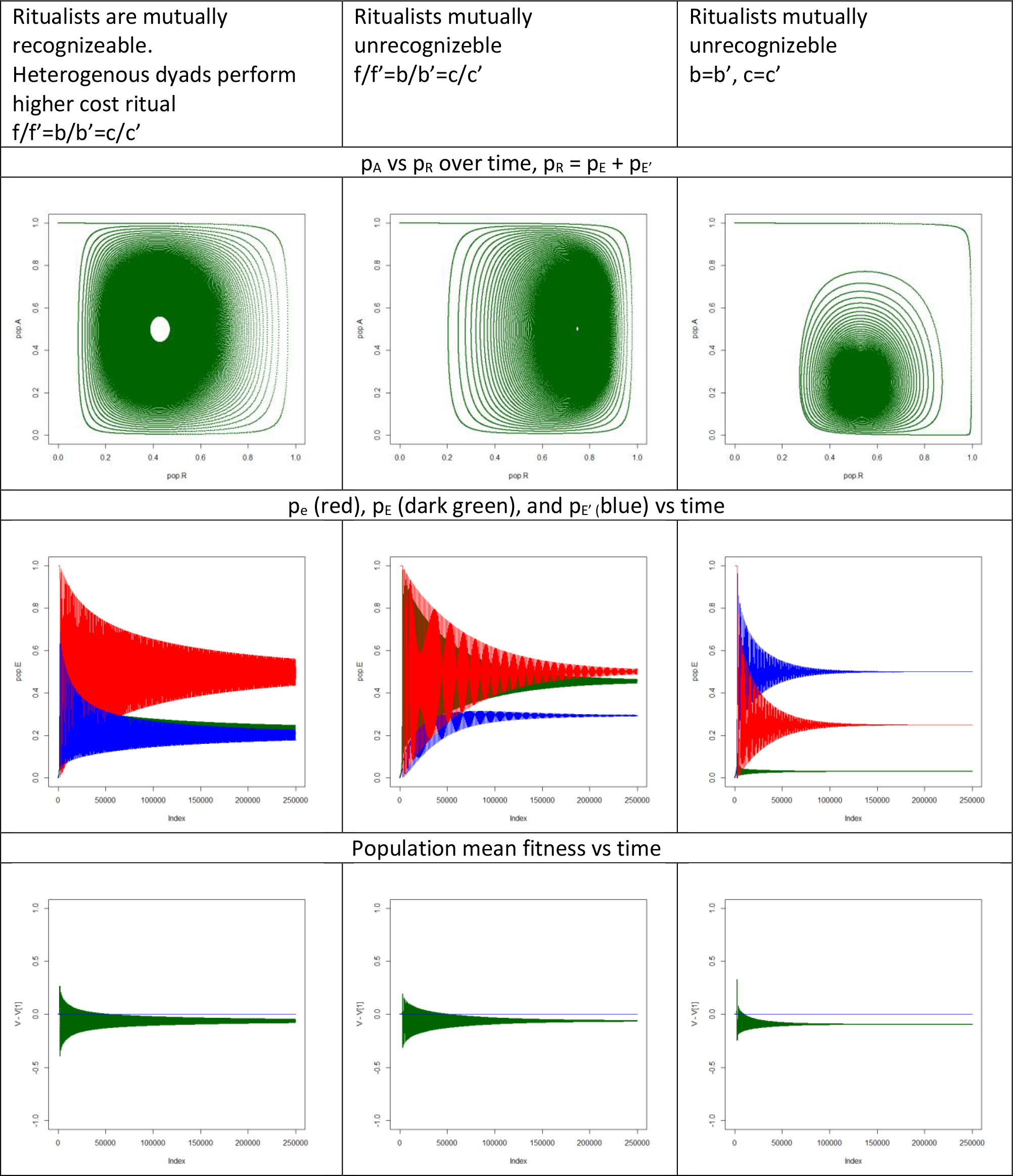
results of simulations for two rituals of different cost: f>f’, p_E,t= 0_ = p_E’,t= 0_ = 0, p_e, t_=0 = 1, p_A,t=0_ = 1

## Appendix 4: Recombination

In the main model of this paper, 100% recombination is assumed. Of course, while recombination rates will be relatively high for genes on separate chromosomes in sexually reproducing species, there is still the possibility of some amount of covariance of traits, and in some cases this can have significant impacts on evolutionary trajectory. To explore this, first I repeated the analysis analytically in a case of zero recombination. Secondly I simulated the evolution with zero recombination. Thirdly, I simulated using a model with varying recombination rates and mutation rates. These simulations use a simpler, more commonly used set of assumptions (Wright-Fisher), rather than the overlapping generations assumptions used in the model referenced in the body of the paper.

## Zero Recombination, analytic solutions

For this model, the fitness functions are the same…

Of course, all 4 types can not coexist at equilibrium since V (ae) never equals V (Ae): these two types can not coexist at equilibrium. Equally obviously each type can exist at equilibrium at fixation, but in all such cases, at least one other variant can invade. AE invades Ae. aE invades AE. Ae or Ae can invade aE. Ae or AE can invade ae. There are no cases where two types can coexist at equilibrium.

There are, however, two situations where 3 types can coexist at equilibrium: absence of Ae or ae.

In the case of absence of ae, the equilibrium is unstable, frequencies given by

- 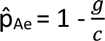
- 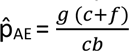
- 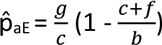

In the case of absence of Ae, the equilibrium is stable (though invadable by Ae), frequencies given by

- 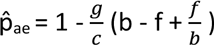
- 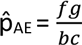
- 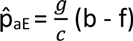

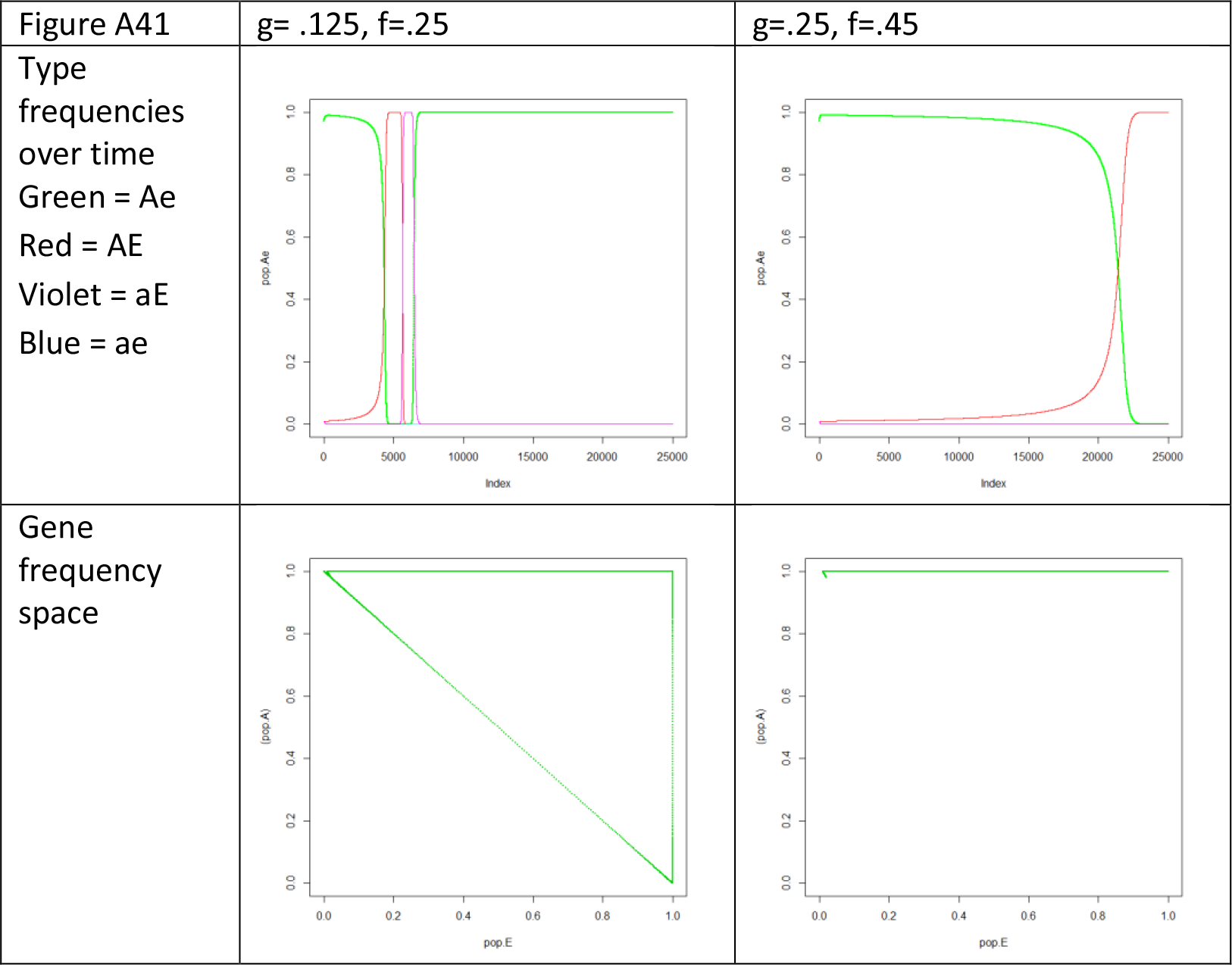

However, this case does not realistically model a sexually reproducing animal, since there will always be some amount of recombination and the effects of mutation may also be significant.

## Zero recombination, simulations

I use simpler Wright Fisher assumptions here of no overlapping generations in an infinite population to model the effects of zero recombination. In each reproductive period a new generation is born in proportion to fitness values of the parent types times the frequency of the parent types. The parents die. This is a more commonly used modeling form, so if the results replicate (to be tested with recombination added back in), then we know that the results are not simply a function of the overlapping generations form of the model.

Without reintroduction of rare types through mutation or recombination, this tends to move the population to relative fixation in either Ae or aE. Figure A1 shows the trajectory of frequencies for a case of each, with a starting population dominated by Ae with trace amounts of the other 3 types (necessary since the main model uses mutation to introduce rare types and mutation is set to zero for this model).

In the formal model, this is again relative fixation, as the structure of the model does not allow for absolute fixation. What has happened here in the first example (g=.125, f=.25) is that the frequency of AE rises, severely reducing the frequency of Ae. Then the frequency of aE rises and the frequency of AE is reduced to over a hundred orders of magnitude lower before Ae has a chance to rise and reduce the population of aE (and thus the frequency dependent pressure against AE). While AE and aE are not eliminated, AE is effectively eliminated (and would be in anything like a real world population characterized by such costs and benefits), which then prevents aE from ever reestablishing itself. In the case of g=.25, f=.45, aE is wiped out in the long time it takes AE to rise, which then prevents Ae from reestablishing itself (via its success in an aE dominated population).

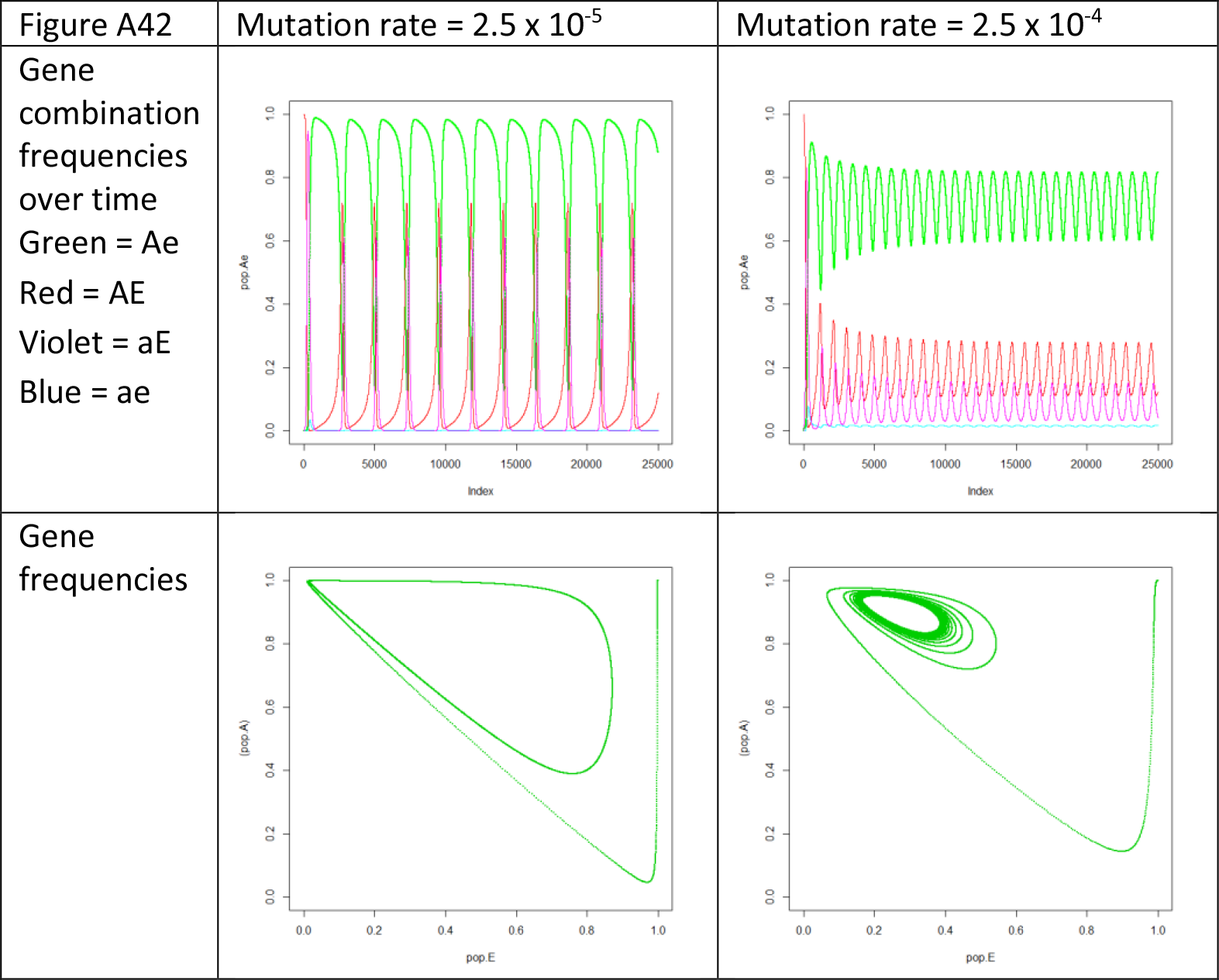

However, with the introduction of a very small amount of mutation (allowing the reintroduction of rare types), the system goes through oscillations similar to the primary model with complete recombination, but with the path of the limit cycle shifted, as illustrated in the Figure A2. As with the primary model, mutation damps the oscillations to a smaller amplitude (see appendix on mutation rates). Where the equilibrium in this case would have been 50/50 for both traits with complete recombination, the system cycles around a point with higher frequencies of A and e, likely largely because of the suppression of ae.

## Partial Recombination, with and without mutation: simulations

The purpose of the modeling exercise is not actually to look at behavior in haploid organisms, but in sexually reproducing diploid animals. The choice of haploid genetics is for simplicity of analysis, to get a feel for the evolutionary dynamics. A sexually reproducing species would have significant recombination rates…50% if on different chromosomes and less but still extant recombination if on the same chromosome. Therefore, I add partial recombination to see how strongly recombination affects the system and how much recombination needs to be present for the system to act like the case of complete recombination. First this is done without mutation and then with mutation to see how they each affect the system.

The results without mutation (Figure A3) show that the system is strongly affected by small amounts of recombination. Instead of going to fixation, even very small amounts of recombination cause the system to move in limit cycles. Moreover, as expected, the system moves to resemble the case of complete recombination with increasing recombination with almost identical dynamics to complete recombination with even small amounts of recombination. Only a little recombination is necessary to mostly eliminate covariance, at least to the point where the remaining covariance is inconsequential for the dynamics. Interestingly, for both very small and for moderate and larger amounts of recombination, the limit cycle has a stable large amplitude, but with small intermediate levels of recombination, the cycles are damped. The recombination in this case is not high enough to change the focal point of the oscillations, but is large enough to act like mutation in terms of reintroducing rare alleles combinations (ae particularly).

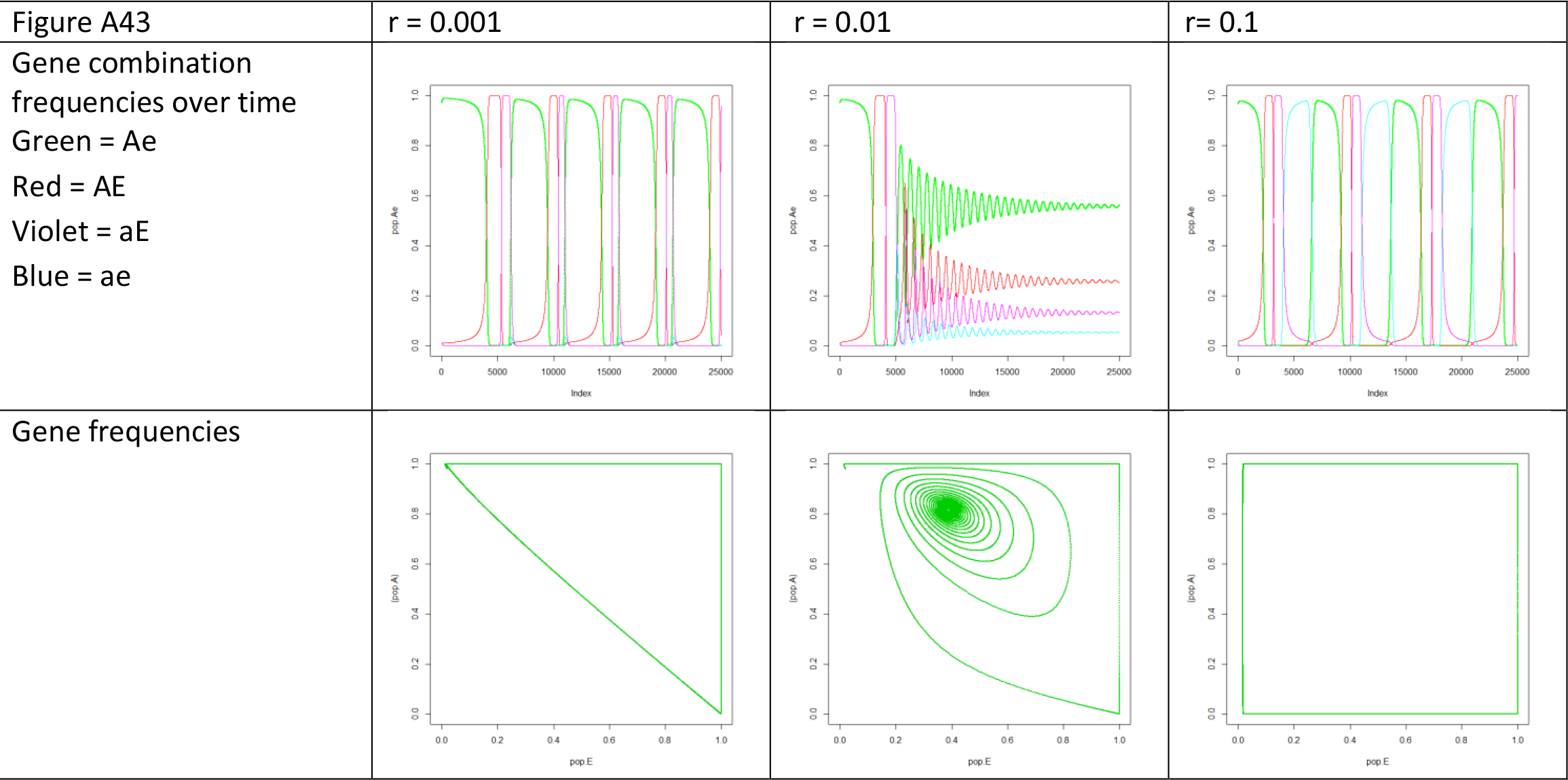

Adding mutation, as expected, dampens the cycles. (Figure A4) This returns the results of the main models analyzed in the body of the paper, including the focal point of the oscillations around the equilibrium frequencies of the model with complete recombination. Again, see the appendix on mutation rates for more exploration of the effects of mutation on the limit cycles. This replication of the results of the main paper with different model assumptions suggests that the simulation results are not just a fluke of the overlapping generations model.

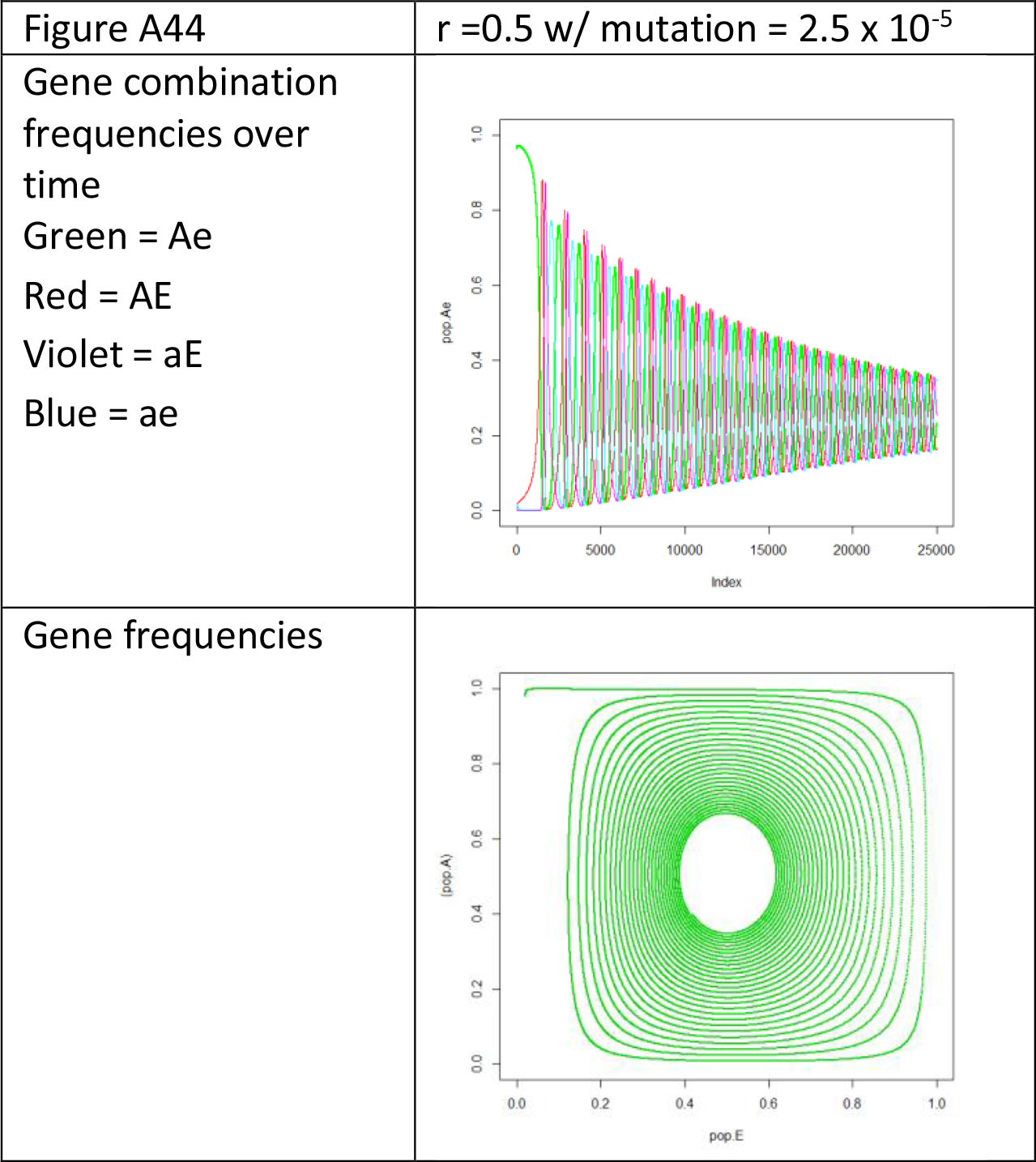

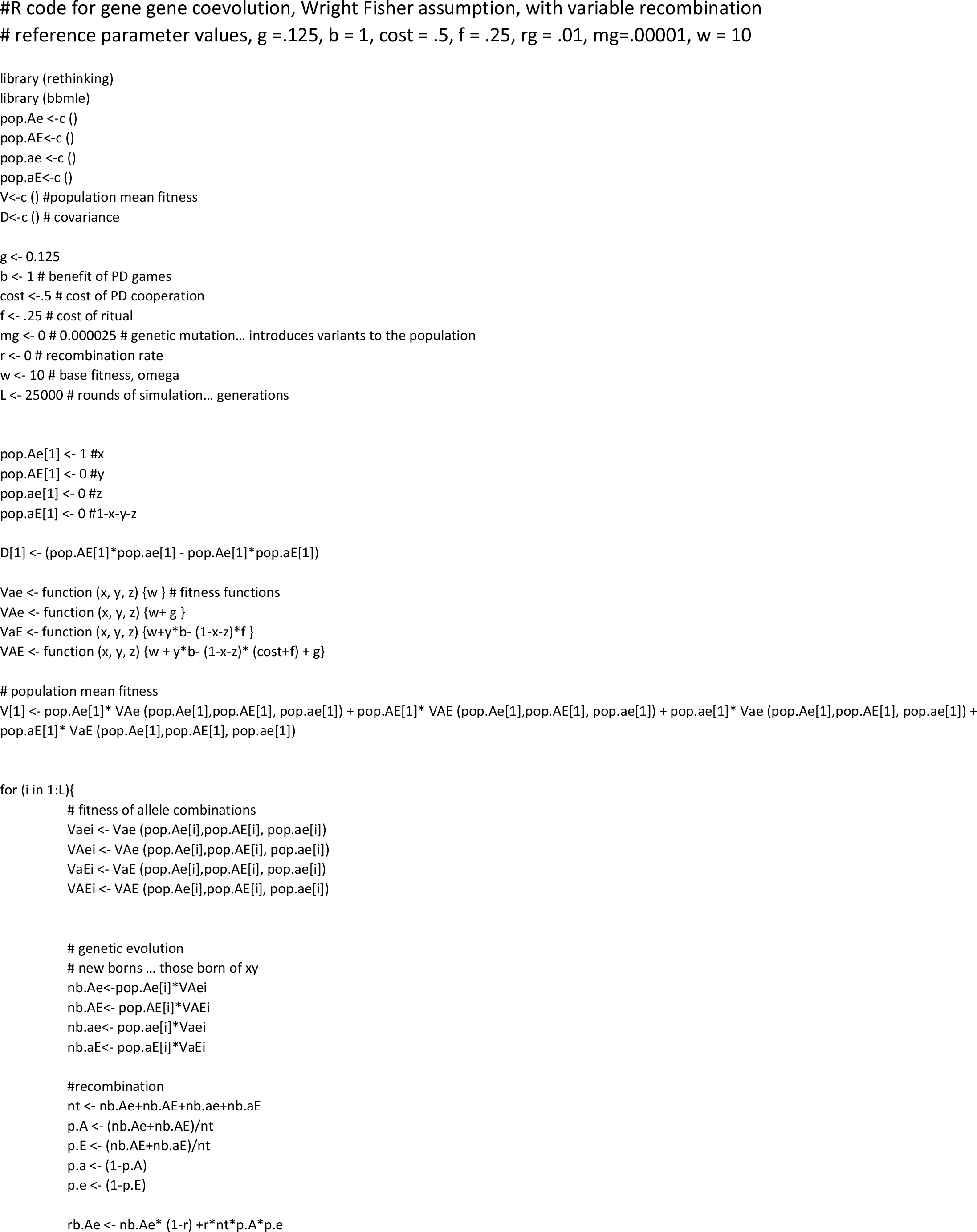

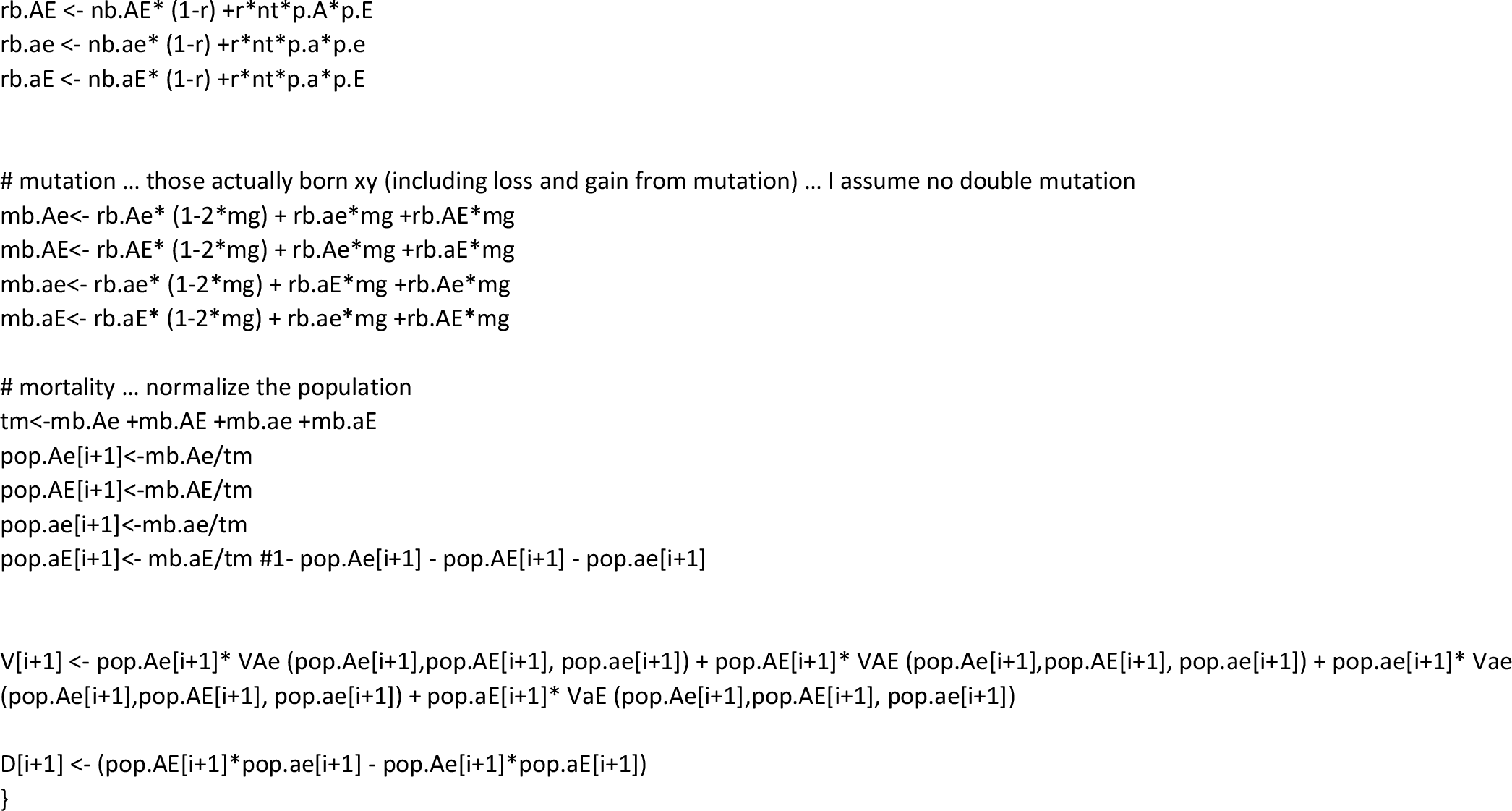

